# Affinity maturation of antibody responses is mediated by differential plasma cell proliferation

**DOI:** 10.1101/2024.11.26.625430

**Authors:** Andrew J. MacLean, Lachlan P. Deimel, Pengcheng Zhou, Mohamed A. ElTanbouly, Julia Merkenschlager, Victor Ramos, Gabriela S. Santos, Thomas Hägglöf, Christian T. Mayer, Brianna Hernandez, Anna Gazumyan, Michel C. Nussenzweig

## Abstract

Increased antibody affinity over time after vaccination, known as affinity maturation, is a prototypical feature of immune responses. Recent studies have shown that a diverse collection of B cells, producing antibodies with a wide spectrum of different affinities, are selected into the plasma cell (PC) pathway. How affinity-permissive selection enables PC affinity maturation remains unknown. Here we report that PC precursors (prePC) expressing high affinity antibodies receive higher levels of T follicular helper (Tfh)-derived help and divide at higher rates than their lower affinity counterparts once they leave the GC. Thus, differential cell division by selected prePCs accounts for how diverse precursors develop into a PC compartment that mediates serological affinity maturation.

## Main text

Increasing antibody affinity over time after vaccination is a prototypical feature of humoral immune responses. Experiments in transgenic mice suggest that in the early germinal center (GC)-independent stages of the immune response, B cells expressing high affinity B cell receptors are preferentially selected into the PC compartment (*1–3*).

Selection within the GC reaction itself is governed by Tfh that recruit B cells and control the degree of B cell clonal expansion (*4*). B cells expressing receptors that bind to and can capture antigen displayed on the surface of follicular dendritic cells in the light zone (LZ) process and present antigens to a limited number of Tfh, in exchange for help signals. Selected LZ B cells initiate a transcriptional program that enables them to move into the dark zone (DZ) where they undergo division and somatic mutation before returning to the LZ to test their newly mutated receptors in subsequent rounds of selection. The number of division cycles and the speed of cell division in the DZ is directly related to the strength of Tfh signals and governed by the level of *Myc* expression (*5–7*). Iterative cycles of DZ division and LZ selection produce GC B cells with increasing affinity for the immunogen over the course of an immune response. However, loss of affinity due to persistent somatic mutation and evolution of the Tfh compartment eventually leads to GC diversification and increasing inclusion of lower affinity B cells (*8–12*).

High affinity cells of GC origin become enriched in the antibody-secreting PC compartment over time after immunization (*13–16*). This observation, in conjunction with the known selection mechanisms within the GC, originally supported a model whereby high affinity cells preferentially undergo PC differentiation. In contrast to these findings, three recent independent studies have shown that in genetically intact animals, diverse collections of GC B cells develop into PC progenitors (*8, 17, 18*). In addition, these studies found little or no enrichment for high-affinity antibody expressing cells among the precursor population, prePCs (*19, 20*), in the GC (*8, 17, 18*). How this population of GC precursors that expresses antibodies with a broad range of affinities gives rise to a PC compartment dominated by cells producing high affinity antibodies is not understood.

### High-affinity cells are over-represented in PC compartment

To examine the relationship between GC B and PCs we tracked the two cell types using fate-mapping S1pr2-CreERT2 R26^lsl-ZSGreen^ mice in which tamoxifen administration permanently labels GC cells and their subsequent products (*21*). Administration of tamoxifen to immunized mice confirmed that this reporter strain labels GC B cells and activated B cells, but not mature PCs (fig. S1A-C). Tamoxifen was administered 5 days after immunization with NP-OVA, and single cell sequencing was performed on labelled GC B and PCs isolated from the draining lymph node on day 14 (Fig. 1A,B). Uniform manifold approximation and projection (UMAP) integrating CITEseq data revealed distinct clusters of GC B and PCs characterized by Fas/CD86 and CD138 surface protein expression respectively (fig. S2).

**Fig 1.**
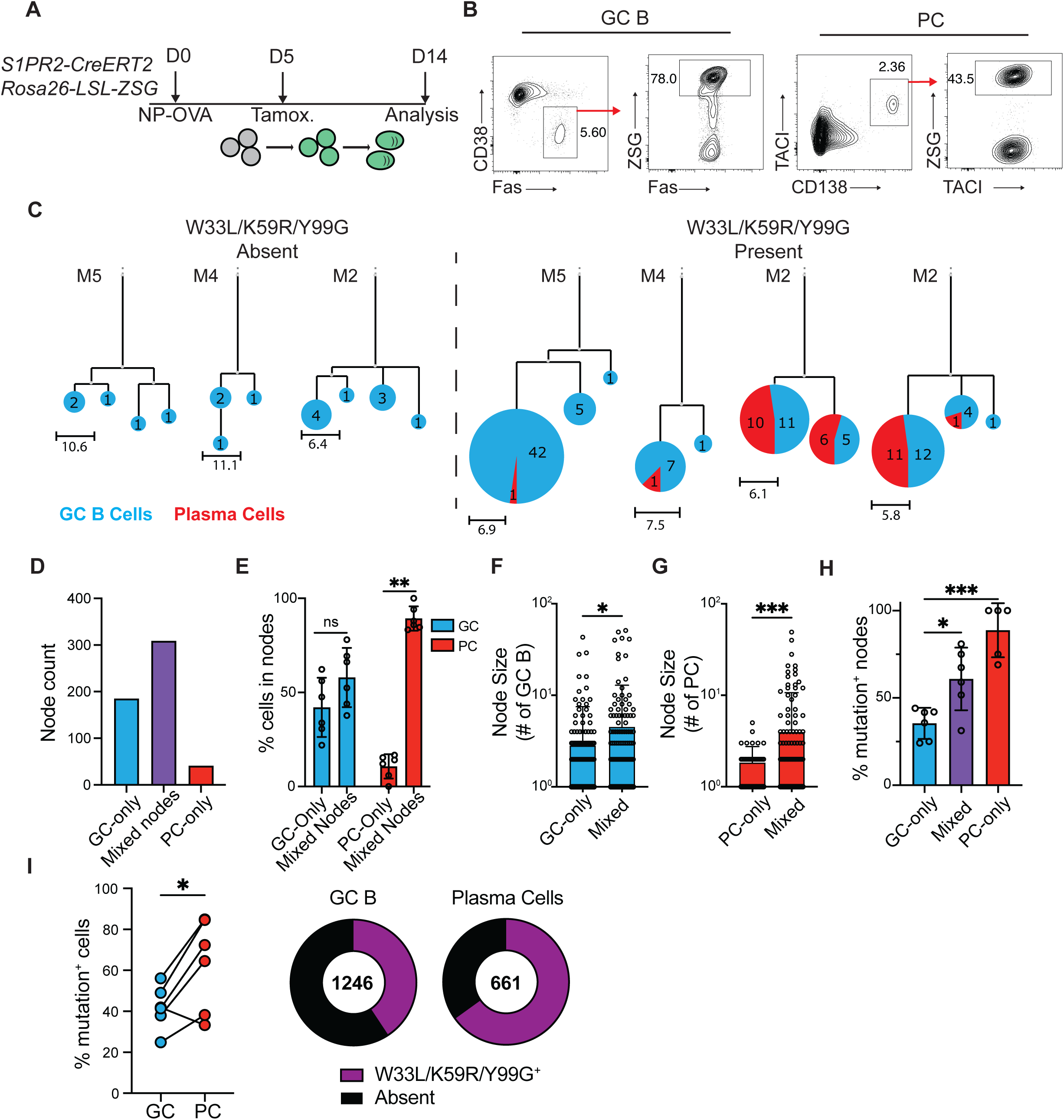
High affinity antibody producing PCs are over-represented relative to contemporaneous GC B cells. (**A**) Experimental layout for (**B-H)**. (**B**), Representative flow cytometry plots showing strategy for isolating popliteal LN (pLN) GC B cells (Pre-gated on live, TACI^-^ CD138^-^ B220^+^; CD38^-^Fas^+^ZSG^+^) and PCs (Pre-gated on live; CD138^+^TACI^+^ZSG^+^). Full gating strategy is displayed in fig. S2A. (**C**) Representative IgH+IgL sequence-based phylogenetic trees highlighting GC B cell (blue) and PC (red) distribution. Expanded clones containing (right) or not containing (left) affinity-enhancing mutations. Each circle represents one node of cells with identical sequences. Scale represents mutational distance observed between related sequences. (**D**) Total numbers of GC-only, PC-only or mixed nodes analyzed. (**E**) Frequency of GC B (blue bars) or PCs (red bars) found within mixed or single cell-type nodes. Each point represents one animal. (**F-G**) Node size; number of individual GC B cells (**F**) or PCs (**G**) in either uniform or mixed nodes. (**H**) Frequency of nodes with affinity-enhancing mutations. Each point represents one mouse, summarizing 51-176 nodes per mouse. (**I**) Frequency of affinity-enhancing mutations within total GC B cells and PCs. Left, summary, each pair of connected points represents GC B cells and PCs isolated from one animal. Right, quantitation of total cells. Numbers in center represent total number of sequences analyzed. * p<0.05, *** p<0.0005. (**F,G**) ANOVA; (**H**) mixed effects analysis; (**I**) paired two-tailed Student’s t-test. Data are pooled from two independent experiments, n=6.

Immunoglobulin sequence analysis showed expanded clones of cells that were further subdivided into nodes containing one or more cells expressing identical antibodies. The combined clonal trees consisted of 535 individual nodes with 1 to >100 members (Fig. 1C). Based on surface staining and gene expression profiles, nodes were divided into those consisting only of GC B cells, both GC B and PCs (“mixed nodes”), or only PCs. Nodes consisting of PCs only were the least abundant, representing <8% of the nodes (38/535 nodes, Fig. 1D). Most PCs were found within mixed nodes that were also larger than either GC- or PC-only nodes (Fig. 1E-G). IGHV1-72 antibody expression is associated with relatively high affinity NP binding (*22*). Notably, compared to GC nodes, nodes containing PCs were enriched for affinity enhancing mutations in IGHV1-72 (W33L/K59R/Y99G (*22, 23*); Fig. 1H). Furthermore, irrespective of node identity, PCs were enriched for affinity-enhancing mutations compared to contemporaneously labelled GC B cells (63±23% PC vs 42±11% GC, p=0.02; Fig. 1I). Thus, although prePC antibody affinity is indistinguishable from contemporaneous GC B cells (*8, 17, 18*), PCs are enriched for high affinity antibodies. Moreover, the presence of identical antibody sequences in large, expanded nodes containing GC and PCs suggests that the mechanisms that govern clonal expansion of the two cell types overlap.

### Differential expansion of high-affinity PCs

Two non-exclusive cellular processes could contribute to the observed enrichment of high-affinity PCs over GC B cells; differential cell death and/or division.

To determine whether cell death contributes to PC affinity maturation, we purified live and dying PCs from draining lymph nodes of NP-OVA-immunized mice using the Rosa26^INDIA/INDIA^ apoptosis reporter mice (*24*). Ig sequence analysis revealed similar abundance of high affinity IGHV1-72 expressing cells in live and dying PC fractions (fig. S3A-D). In addition, most Ig sequences found among apoptotic PC clones were also present in the live fraction (fig. S3C). Thus, differential cell death does not appear to drive the accumulation of high affinity PCs over time.

To determine whether high affinity PCs have a proliferative advantage we tracked cell division using Vav^tTa^ Col1A1-tetO-histone H2B^mCherry^ reporter mice (H2B-mCherry) (*25*). Under steady state conditions, tTA is expressed in hematopoietic cells and induces high levels of the histone H2B–mCherry fusion protein expression. Administration of doxycycline represses histone H2B– mCherry synthesis, and therefore mCherry fluorescence dilutes in proportion to cell division (*25*). H2B-mCherry mice were immunized with NP-OVA and administered doxycycline 10 days after immunization. PCs were isolated from draining lymph nodes 2 days later on the basis of their high or low levels of division (mCherry^lo^ and mCherry^hi^ PCs, respectively; Fig. 2A,B). mCherry^lo^ PCs that had undergone greater levels of division showed reduced species richness and larger clonal families when compared to their non-dividing counterparts, indicating rapid clonal expansion in this population (Fig. 2C-E). Notably, there was an enrichment of PC expressing high-affinity IGHV1-72 among the more divided PCs (75.1±14% mCherry^lo^ vs 37.1±12% mCherry^hi^; p=0.0001; Fig. 2F,G). At this early timepoint the W33L/K59R/Y99G affinity enhancing mutations were largely absent from both IgHV1-72^+^ PCs populations, but there was a trend towards enrichment in the more divided mCherry^lo^ group (Fig. 2H, p=0.08). These findings were confirmed using Blimp1-CreERT2 R26^lsl-ZSGreen^ mice, a reporter strain which selectively labels differentiated PCs (fig. S4A-B). Blimp1-CreERT2 R26^lsl-ZSGreen^ mice were combined with H2B-mCherry mice to enable tamoxifen-mediated ZSG-labelling of a synchronized cohort of PCs which can be division-tracked by mCherry dilution. Isolation of mCherry^Hi^ and mCherry^Lo^ ZSG^+^ PCs revealed that amongst mature PCs, the more proliferative cells were enriched for NP bait binding, and expression of high-affinity Ig gene rearrangements (fig. S4C-H). These data indicate that higher affinity B cell receptor expression is associated with differential division at the mature PC stage.

**Fig 2.**
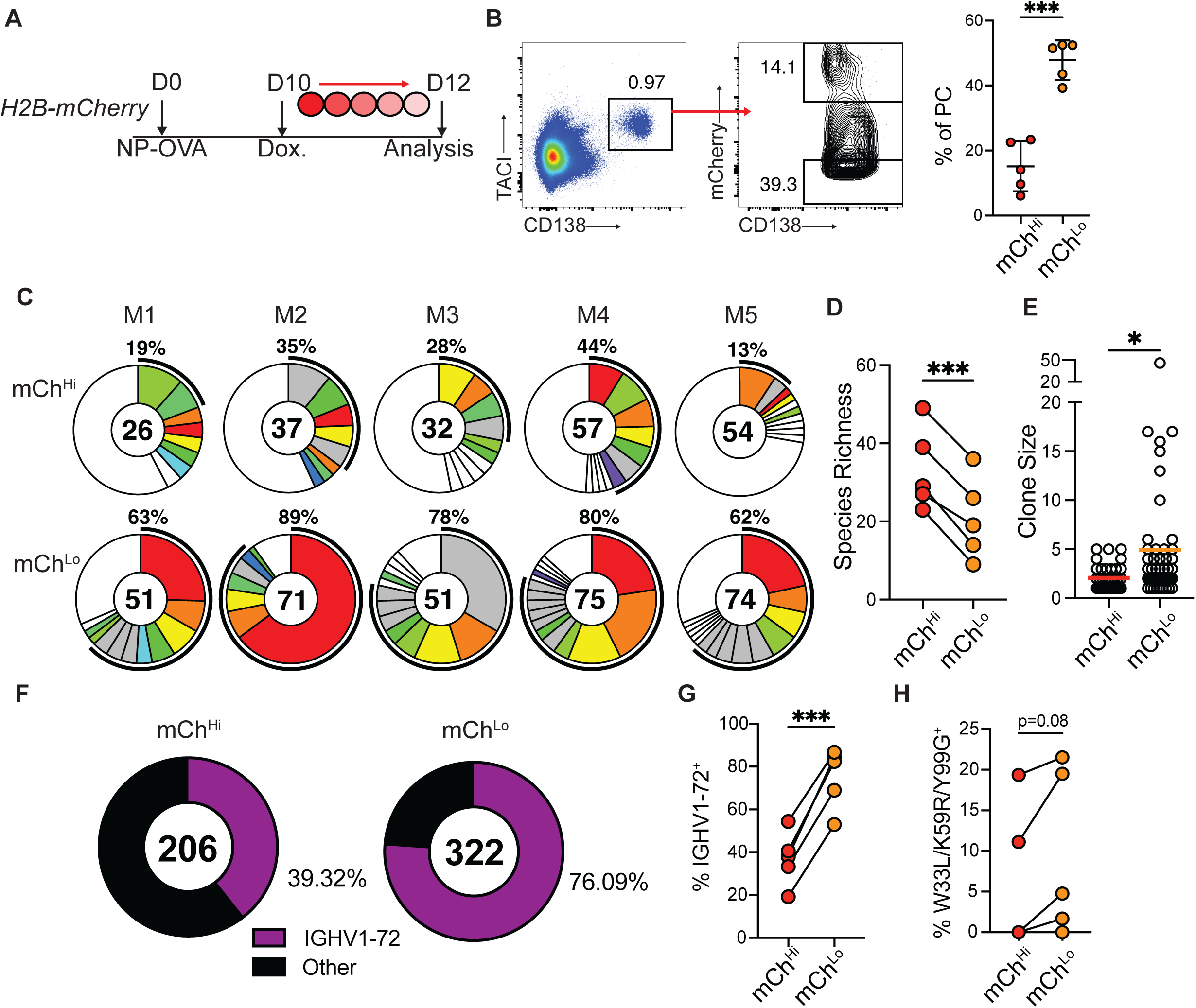
High affinity antibody producing PCs are more proliferative. (**A**) Experimental layout for (**B-F**). (**B**) Left, flow cytometry profile showing TACI^+^CD138^+^ PCs and gating for mCh^hi^ and mCh^lo^ cells from pLNs. Right, mCh^lo^ and mCh^hi^ PC frequency. Each point represents one mouse. Graphs display mean±SD. (**C**) Clonal distribution of paired Ig sequences (IgH+IgK/IgL) among mCh^hi^ and mCh^lo^ PCs isolated from each mouse. Colored segments represent expanded clones and singlets are represented by white segments. The number in the center represents the number of sequences analyzed per population. The outer segment annotation denotes the percentage of cells that are members of expanded clones. (**D**), Chao1 species richness quantification. Each pair of points represents one mouse. (**E**) Clone size. Each point represents one clone. (**F**) Frequency of mCh^hi^ or mCh^lo^ PCs bearing high-affinity IGHV1-72 BCRs. Number in center represents total number of sequences. (**G**) Summary of (**F**) each pair of points represents one mouse. (**H**) Frequency of high affinity mutation containing sequences among IGHV1-72^+^ expressing mCh^Hi^ or mCh^Lo^ PCs. * p<0.05, *** p<0.0005. (**B,D,G,H**) paired two-tailed Student’s t-test; (**E**) unpaired Student’s t-test. Data are pooled from 2 independent experiments, n=5.

To validate these findings using vaccine antigens, we immunized H2B-mCherry mice with either SARS-CoV2 receptor binding domain (RBD) or combined tetanus and diphtheria toxoid antigens (Tenivac^®^) and administered doxycycline on day 10 after vaccination. Antigen binding was used as a surrogate for higher affinity antigen binding cells (fig. S5A,B). Flow cytometric analysis revealed that RBD or tetanus and diphtheria toxoid binding cells were enriched and showed higher mean fluorescence among the divided mCherry^lo^ compared to mCherry^hi^ cells, even when normalized for surface Ig expression levels (fig. S5C-H). The combined data set indicates that high affinity plasma cells underwent greater levels of clonal expansion than lower affinity PCs irrespective of the composition of the administered immunogen or adjuvant formulation.

### PC selection outside of the GC

To examine the site of affinity-driven PC clonal expansion, S1pr2-CreERT2 R26^lsl-ZSGreen^ mice were immunized with NP-OVA, treated with tamoxifen on day 8 and then given anti-CD40L or isotype control antibodies on day 10 and 12 after immunization. In this setting, tamoxifen will label GC products and anti-CD40L will terminate the GC reaction 2 days later, allowing a cohort of PCs generated between days 8-10 to be tracked. Draining lymph nodes were examined by confocal microscopy on day 16 after immunization (Fig. 3A,B, and fig. S6A).

**Fig 3.**
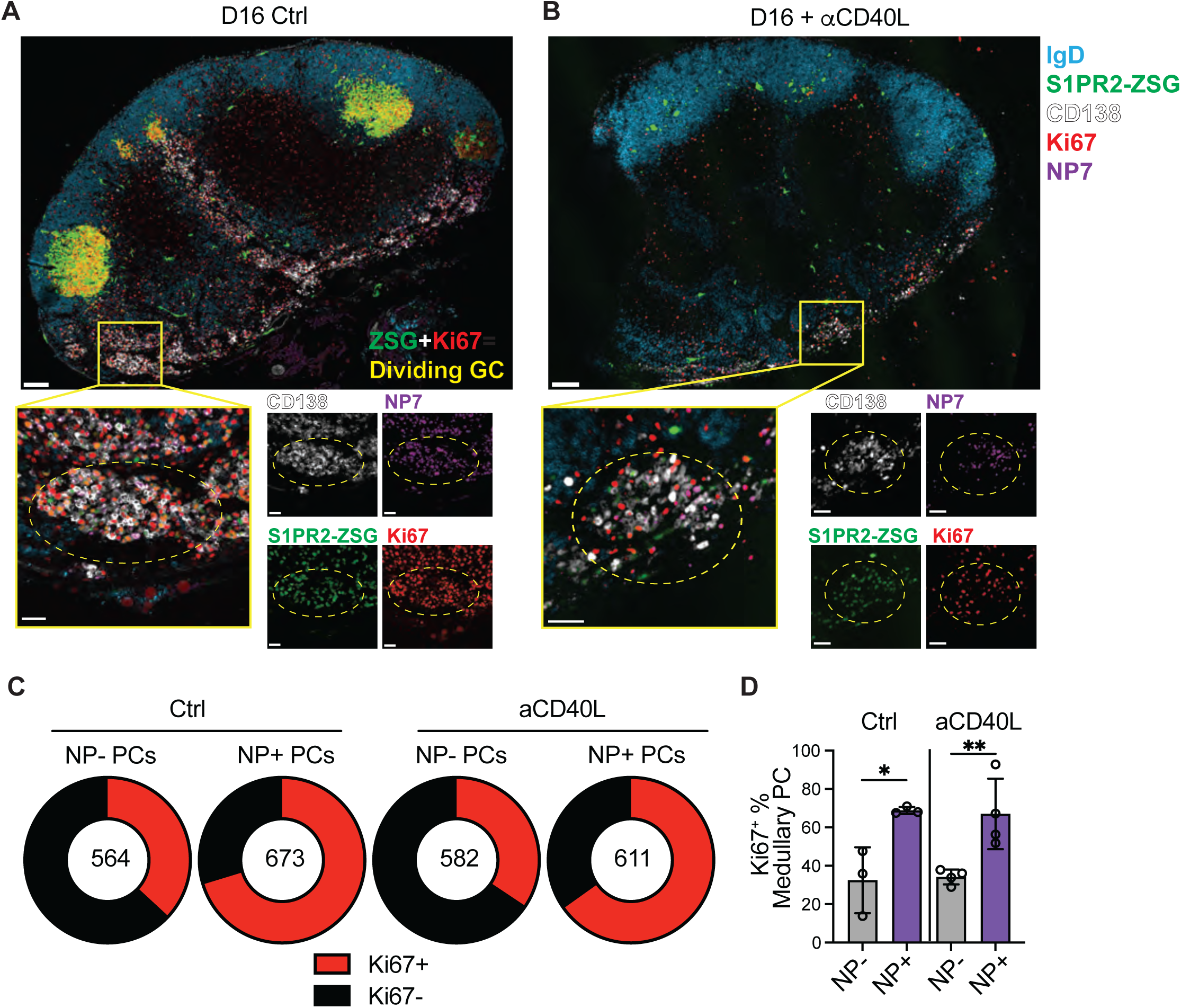
Clonal evolution of the PC repertoire. (**A-B**), Confocal microscopy images of pLNs isolated from S1pr2-CreERT2 R26^lsl-ZSGreen^ mice immunized with NP-OVA, treated with tamoxifen on D8 and anti-CD40L or isotype control on D10 and D12. Large tiles show overview of pLN architecture. Inset boxes show individual clusters of dividing Ki67^+^ NP^+^ PCs that are highlighted by dashed circles. Individual channels are shown beside merged images. In large tiles, scale bar=100um. In inset boxes, scale bar=30um. (**C**) Quantitation of fraction of medullary PCs which are Ki67^+^ amongst NP^+^ and NP^-^ populations, as quantified from confocal images as in (**A**). The number in the center represents the number of cells analyzed per population. (**D**) Summary of (**C**); each point represents one mouse. Graph displays mean±SD. *p<0.05, **p<0.005; ordinary one-way ANOVA. Data are representative of three experiments, n=3-4 per group.

Large GCs in which many of the cells were actively dividing (as determined by positive staining for Ki67) were observed in control mice that did not receive anti-CD40L. In addition, we found discrete clusters of dividing NP-binding PCs (CD138^+^ZSG^+^Ki67^+^NP^+^) which were particularly abundant in the LN medulla. Anti-CD40L treatment aborted the GC reaction (Fig. 3A-B, and fig. S6A-B). In contrast, the clusters of proliferating NP-specific PCs persisted (Fig. 3A-B). Quantitation of medullary PC division by Ki67 staining revealed a higher proportion of dividing NP-binding than non-binding PCs. Moreover, this feature was retained upon anti-CD40L treatment and GC depletion (Fig. 3C-D). Thus, dividing antigen-specific PCs were observed in medullary foci outside of the anatomical confines of the GC, and these foci were maintained in the absence of active GCs during the observation period.

To determine whether the observed enrichment of high affinity PCs continued in a post-GC compartment, we isolated plasma cells from anti-CD40L treated S1pr2-CreERT2 R26^lsl-ZSGreen^ mice. The mice also received the drug FTY720, to inhibit S1PR1-driven egress signals and to prevent PCs from leaving the node (*26*). Treatment with anti-CD40L effectively depleted draining lymph nodes of GCs and activated B cells within 2 days (Fig. 3A,B, and fig. S6A-D). Immunoglobulin sequence analysis on day 10 showed that 67% of labelled PCs were IGHV1-72^+^, and that by day 16 this increased to 83% under control conditions (p=0.037, Fig. 4A-E). Notably, the PC compartment was equally enriched for high affinity clones on day 16 in mice treated with anti-CD40L, under conditions that depleted GCs (Fig. 4C-E). In addition, the PC compartment in CD40L-treated mice displayed the same characteristic reduction in species richness that was indicative of clonal expansion seen in control mice (Fig. 4B,F). Finally, labeled PCs obtained from anti-CD40L-treated mice displayed a reduced mutational load, suggesting that they were derived from GC precursors that exited the GC at a time when mutations were less abundant (Fig. 4G).

**Fig 4.**
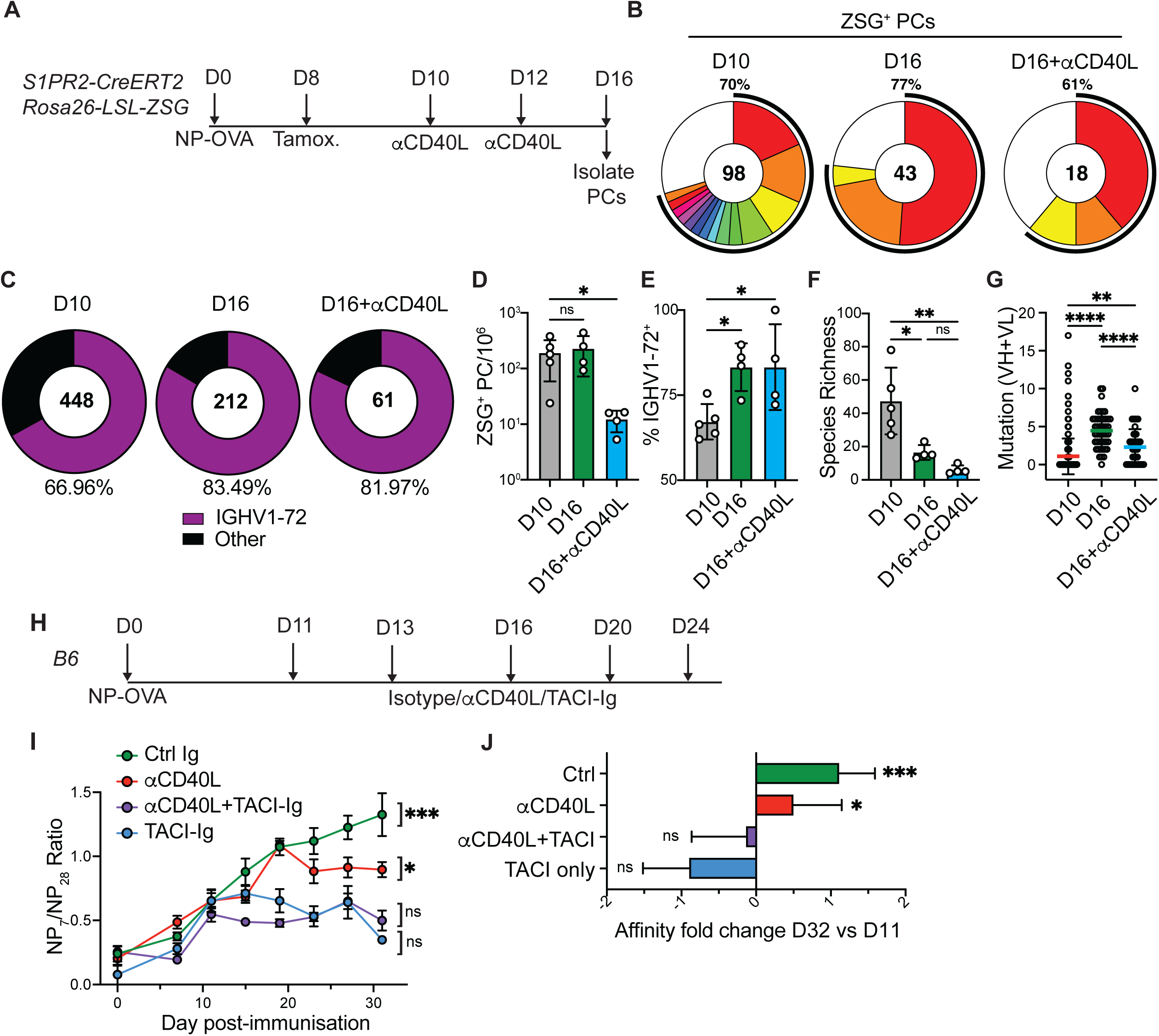
Serological affinity maturation. (**A**) Experimental layout for (**B-G**). (**B**) Clonal distribution of paired Ig sequences from representative mice. Colored segments represent expanded clones and singlets are represented by white segments. The number in the center represents the number of sequences analyzed per population. The outer segment annotation denotes the percentage of cells that are members of expanded clones. (**C**) Frequency of ZSG^+^ PCs expressing IGHV1-72. The number in the center represents the number of sequences analyzed. (**D**) Number of ZSG^+^ PCs in each experimental group. (**E**) Summary of IGHV1-72 frequency presented in (**C**). (**F**) Chao1 species richness. In D-F, each point represents one mouse, n=4 -5 per condition. (**G**) Number of VH+VL mutations in paired Ig sequences. Each point represents one cell. (**H**), Experimental layout for (**I-J**). (**I**) Ratio of NP_7_/NP_28_-binding IgG in serum measured by ELISA. (**J**) Fold change in affinity (NP_7_/NP_28_ ratio) at D32 vs D11. Data are presented as mean±SD. Data are representative of two experiments, n=5-10 per group. * p<0.05, ** p<0.005. **** p<0.0001. (**D-F**) ordinary one-way ANOVA; (**G**) Kruskal-Wallis test. (**I-J**), mixed effects analysis. For full statistical comparisons see Supplementary Fig.4E.

Serological affinity maturation is typified by a rapid increase in the affinity of circulating antibodies (*27*). We hypothesized that the continued preferential expansion of high affinity PCs after GC-depletion (Fig. 4A-G) would lead to measurable improvements of serum anti-NP affinity. To determine whether PCs in anti-CD40L treated mice continue to support affinity maturation after depletion of GC and activated B cells, we measured the serum antibody binding to bovine serum albumin derivatized with either 7 or 28 molecules of NP (Fig. 4H and fig. S6A-E). Control mice displayed a marked increase in NP_7_/NP_28_ IgG binding between days 12 and 32 indicating a rapid increase in anti-NP affinity that was aborted by depleting PCs with TACI-Ig (Fig. 4I). Notably, and consistent with the above sequencing data, the serum antibody response continued to undergo affinity maturation for several weeks after anti-CD40L treatment (Fig. 4I,J).

We conclude that serological affinity maturation can proceed in the absence of continued output from the GC reaction, and it is suppressed by plasma cell depletion. These findings suggest that post-export from the germinal center, newly generated PCs undergo affinity-based expansion in a manner that contributes to increasing serum antibody affinity.

### PC division is proportional to the strength of T cell help

GC B cells in the LZ of the GC structure compete for limited help signals from T follicular helper (Tfh) cells, a process known as positive selection. The amount of help received by a B cell is directly proportional to the amount of antigen captured and presented by the B cell (*25, 28*). Positively selected GC B cells upregulate expression of the transcription factor *Myc* in proportion to the strength of T cell signals and migrate to the DZ where they undergo cycles of Tfh independent inertial cell division in proportion to the amount of *Myc* expression (*5, 25, 29*).

To determine whether a similar process regulated the relative amount of PC expansion, we delivered graded doses of the antigen ovalbumin (OVA) to ongoing GCs using chimeric anti-DEC205-OVA antibodies (*28*) (Fig. 5A). Protein antigens loaded onto anti-DEC205 are delivered to DEC205^+^ GC B cells, supplying a B cell receptor-independent supply of antigen which can be processed and presented to Tfh cells (*28*). As expected, GC B cells expanded in direct proportion to the amount of antigen delivered (Fig. 5B,C). Antigen delivery by anti-DEC-205-OVA also produced a rapid increase in the frequency and numbers of prePCs (B220^hi^ CD38^-^ Fas^+^ CD138^+^; Fig. 5B,D). To determine whether prePCs in the LZ of the GC also expressed *Myc* in proportion to antigen delivery by DEC-205, we purified these cells 24 hours after graded doses of anti-DEC-205-OVA injection. Consistent with their commitment to the plasma cell fate these cells displayed elevated levels of *Irf4*, a transcription factor which promotes PC differentiation, and lower expression levels of the antigen receptor signaling-induced factor *Nr4a1* than other LZ GC B (fig. S7A,B). Notably they showed increases in *Myc* that were directly proportional to the amount of antigen delivered (Fig. 5E). *Tfap4*, a transcription factor downstream of c-Myc, also showed a similar induction pattern, although the expression of this factor plateaued at lower antigen concentrations (Fig. 5F). In addition, PrePC proliferation, as measured by Ki67 expression, was proportional to the amount of antigen supplied (fig. S7C,D).

**Fig 5.**
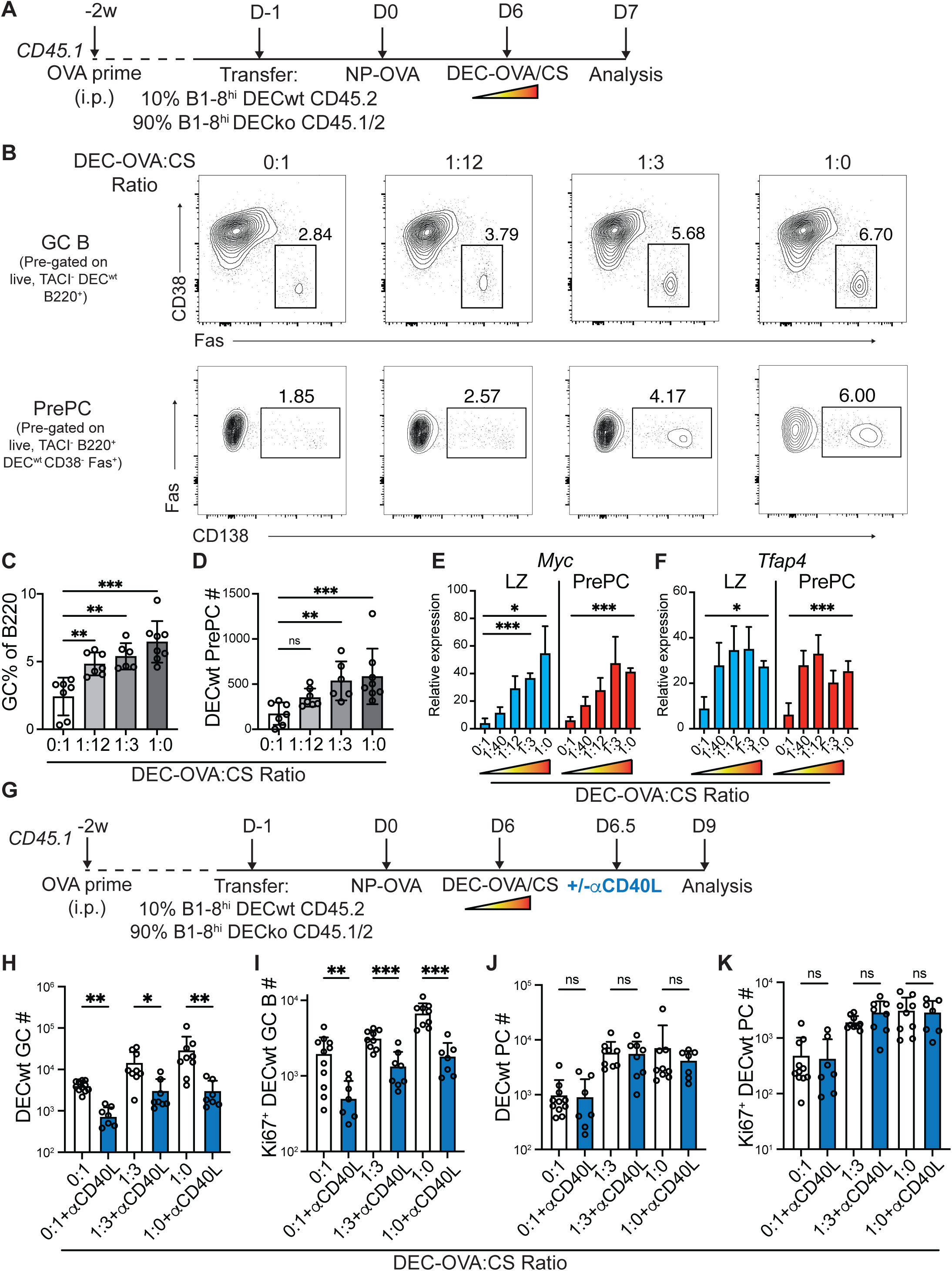
PC proliferation is proportional to signal strength. (**A**) Experimental outline for (**A-F**). (**B**) Representative flow cytometry plots showing GC B cells (B220^+^CD38^-^Fas^+^, top) and prePCs (B220^+^CD38^-^Fas^+^CD138^+^, bottom), after treatment with different ratios of anti-DEC-OVA:CS. (**C-D**) Quantitation of DEC^wt^ GC B cells (**C**) and prePCs (**D**) 24h after anti-DEC treatment. (**E-F**) qPCR of sorted DEC^wt^ GC LZ (CD86^+^CXCR4^-^ GC B) and prePC (as above) showing GAPDH-normalized relative expression for *Myc* (**E**) and *Tfap4* (**F**). Data in (**A-D**) and (**E-F**) are representative of 3 independent experiments, n=6-8 per group. (**G**) Experimental outline for (**H-K**). (**H-I**) Quantitation of DEC^WT^ GC B cells (**H**) and Ki67^+^ DEC^WT^ GC B cells (**I**). (**J-K**) Quantitation of DEC^WT^ PCs (**J**) and Ki67^+^ DEC^WT^ PCs (**K**). Data are presented as mean±SD. Data in (**H-K**) are representative of 4 independent experiments. * p<0.05, ** p<0.005. (**C**) ANOVA; (**D, H-K**) Kruskal-Wallis test.

To test whether Tfh help in the LZ could support continued PC expansion, anti-CD40L was administered 12 hours after anti-DEC-205-OVA. In this setting GC B cells acquired high levels of cognate antigen and interact with Tfh for ∼12h, after which subsequent T cell help was inhibited by CD40L blockade as evidenced by GC collapse (Fig. 5G, Fig. 3A,B, and fig. S6A-B). Treatment with anti-CD40L 12h after anti-DEC205-OVA injection prevented further GC B cell expansion and blocked additional Ki67 expression in GC B including prePCs (Fig. 5H,I, and fig. S7E,F). In contrast, mature PCs continued to expand and divide in the days following anti-CD40L-treatment in direct proportion to the amount of antigen captured (Fig. 5J,K). The data indicate that after a short pulse of T cell help, developing prePCs upregulate *Myc* and undergo cycles of cell division in the absence of continued T cell signals.

Other signals are known to enhance the magnitude of the PC response (*14*). For example, PCs are highly sensitive to the levels of the cytokine IL-21 (*30, 31*), and IL-21-producing T cells have been reported to localize in foci outside of the B cell follicle (*32*). To test whether IL-21 could support the expansion of committed PCs, we fate-mapped PCs on day 8 using Blimp1-CreERT2 reporter mice and subsequently treated with anti-CD40L or anti-IL21R from days 10-12. PC division was assessed by Ki67 staining. As expected, analysis of ZSG-labelled PCs revealed that continued CD40L signals were not required to sustain PC proliferation (fig. S8A-C). In contrast, partial blockade of IL-21R led to reduced proliferation of mature ZSG^+^ PCs, indicating that IL-21 supports the post-GC expansion of plasma cells.

## Discussion

Current models of PC differentiation suggest that B cell receptor affinity is deterministic of cell fate (*14, 15, 27*). Our experiments indicated that while a heterogeneous collection of prePCs, including those expressing low affinity receptors, developed into PCs they subsequently underwent differential affinity dependent division. As a result, PCs expressing higher affinity antibodies contributed disproportionately to serum antibodies, promoting affinity maturation.

Antibody responses develop in two stages, a GC independent early phase during which B cells rapidly develop into dividing plasmablasts that produce relatively lower affinity antibodies and a second GC dependent phase that produces higher affinity antibodies responsible for serologic affinity maturation. In transgenic mice that carry high affinity antibodies, the early GC independent rapid burst of PC development and expansion is affinity dependent (*1, 2*). In contrast, under physiologic circumstances this early PC response produces primarily germline encoded antibodies with relatively lower affinity than those produced in GCs (*15, 33, 34*). In animals with a polyclonal B cell repertoire, GCs enable B cell clonal expansion and antibody hypermutation, both of which are essential for antibody affinity maturation (*27, 35, 36*). Our experiments elucidate the mechanisms by which GC-dependent PC selection enables affinity maturation in polyclonal immune responses.

The GC LZ also contains prePCs that express IRF4, CD138, variable levels of Myc, and share many of the transcriptional features of LZ B cells selected for DZ re-entry (*8, 14*). Like DZ B cells, PC descendants of prePCs undergo rapid cell division in GC adjacent sites but they do not undergo additional somatic mutation and so retain antibody specificity (*37*). Our experiments show that the amount of clonal expansion by newly exported PCs is directly proportional to affinity and the strength of historic selection signals these cells received in the LZ, providing a mechanistic explanation for how PC selection is regulated in relation to affinity. Notably, our data is entirely consistent with the observation that the PC pool tends to be more clonal than contemporary GC B or pre-PCs from which they develop (*8, 17*). In addition to PCs in the LNs, long-lived PCs in the bone marrow are also enriched for high affinity antibody producing cells (*38, 39*). Our data suggests that this phenomenon is likely due to over-representation of high affinity PCs among dividing plasmablasts that subsequently seed the bone marrow.

In summary, post-GC PC clonal expansion in proportion to historical Tfh signals and *Myc* expression parallels Myc-regulated GC B cell clonal expansion, thereby enabling rapid affinity maturation. The proposed model does not preclude permissive selection of GC cells with diverse affinities into the PC compartment (*8, 17, 18*). Instead, our findings help resolve the apparent contradiction between affinity permissive selection into the PC compartment and rapid affinity maturation.

## Acknowledgements

We thank T. Kurosaki for providing S1pr2-CreERT2 mice, T. Eisenreich for mouse colony management, K. Gordon for cell sorting, Anna Minh Newen for INDIA mice immunizations and all members of the Nussenzweig laboratory for helpful discussion.

## Funding

This work was supported in part by NIH grant 5R37 AI037526 and NIH Center for HIV/AIDS Vaccine Immunology and Immunogen Discovery (CHAVID) 1UM1AI144462-01 to M.C.N and the Stavros Niarchos Foundation Institute for Global Infectious Disease Research. J.M. is a Branco Weiss fellow.

M.C.Nussenzweig is a Howard Hughes Medical Institute (HHMI) Investigator.

## Author Contributions

A.J.M and M.C.N conceived the study, designed experiments, interpreted data and wrote the manuscript. A.J.M, L.P.D, P.Z and J.M. designed and performed experiments. V.R. and G.S.S performed bioinformatic data analysis. M.A.E. designed and generated the Blimp1-CreERT2 mouse strain. A.G., T.H., M.A.E., C.T.M. and B.H. designed and produced critical reagents. A.J.M, L.P.D. ,P.Z., J.M., V.R., G.S.S., T.H., M.A.E., C.T.M., B.H., A.G., and M.C.N contributed to editing the manuscript.

## Data and materials availability

All data needed to evaluate the conclusions in the paper are available in the main text and supplementary materials. Sequencing datasets have been deposited in NCBI’s Gene Expression Omnibus, and are available through GEO accession number GSE282284. Correspondence and requests for materials such as mouse strains should be addressed to A.J.M and M.C.N., under a material agreement with The Rockefeller University.

## License Details

This article is subject to HHMI’s Open Access to Publications policy. HHMI lab heads have previously granted a nonexclusive CC BY 4.0 license to the public and a sublicensable license to HHMI in their research articles. Pursuant to those licenses, the Author Accepted Manuscript (AAM) of this article can be made freely available under a CC BY 4.0 license immediately upon publication.

## Materials and Methods

### Mice

S1pr2-CreERT2 mice were kindly provided by T. Kurosaki (*21*). B1-8^hi^, mCherry and DEC^ko^ were as described (*13, 25, 28*). R26^lsl-ZSGreen^ and C57Bl/6J mice were purchased from Jackson Laboratories. R26^INDIA/INDIA^ were kindly provided by C. Mayer (*24*). Blimp1-CreERT2 mice were generated by insertion of a tamoxifen inducible Cre into the 3’ untranslated region of the Prdm1 (Blimp1) gene (Blimp1-CreERT2) linked by a P2A sequence. To verify the specificity of the Blimp1-CreERT2 driver we combined it with the R26^lsl-ZSGreen^ indicator allele. Male mice aged 6-12 weeks were used in all experiments. Animals were housed at The Rockefeller University Comparative Bioscience Center, and all animal procedures were performed following protocols approved by the Rockefeller University Institutional Animal Care and Use Committee. Animals were housed at an ambient temperature of 22C and a humidity of 30–70% under a 12h–12h light– dark cycle with ad libitum access to food and water. For experimental endpoints, mice were euthanized by cervical dislocation.

### Immunizations and tamoxifen treatment

NP-OVA immunizations were performed by subcutaneous footpad injection of 12.5ug NP_16_-OVA (Biosearch Technologies) in 33% alhydrogel (Invivogen). Recombinant SARS-CoV2 RBD was produced as described (*40*). For RBD immunization, 5ug RBD was administered per footpad, in 33% alhydrogel. 50ul of Tenivac^®^ (tetanus and diphtheria toxoids, adsorbed; Sanofi Pasteur) was administered per footpad. For tamoxifen administration, each mouse received one dose of 12mg tamoxifen (Sigma-Aldrich, T5648) prepared in corn oil by oral gavage at the indicated timepoint.

### Antibody infusion and FTY720 administration

For GC experiments 500ug of anti-CD40L (Clone MR1; BioXCell) was administered intravenously on D10 and 100ug on D12 after immunization. For longitudinal serological experiments 100ug anti-CD40L was also given on D16, D20 and D24 to maintain depletion.

For partial IL-21R blocking, 50ug of anti-IL-21R (Clone 4A9, BioXCell), was administered subcutaneously on D10.

For PC depletion, TACI-Ig was produced as described (*41*), with the following differences. Briefly, an expression vector for TACI-Ig fusion protein was generated by conjoining DNA sequences encoding the pre-pro signal sequence from human tissue plasminogen activator, the extracellular domain of mouse TACI (aa2–82), and a mutated H chain Fc region from the C57BL/6 mouse IgG2c (lacking CH1). The L235E, E318A, K320A, and K322A aa substitutions were introduced based on codon usage into BL/6 IgG2c Fc as detailed (*41*). TACI-Ig was produced in Expi293F (Thermo Fisher Scientific). 100ug TACI-Ig was administered intravenously on D10, D12, D16, D20 and D24.

To prevent PC egress from LNs, 1mg/kg FTY720 (Selleck Chemicals) was administered intravenously on D10 and D14 after immunization as indicated.

Anti-DEC205-CS and anti-DEC205-OVA were produced as described (*42*). Mixtures of anti-DEC205-OVA and -CS were administered at the indicated ratios, with a total of 5ug administered to each footpad, on D6 after immunization. For DEC experiments, a cell transfer approach was utilized to allow targeted antigen delivery to a fraction of GC B cells (90% DECko/10%DECwt B cells). Previous experiments in our lab have shown this approach to maintain the competitive nature of the GC reaction, and allow the help to be focused on a small proportion of responding cells (*28*).

### Bait preparation

Avi-tagged recombinant SARS-CoV2 Wuhan Hu-1 RBD (*40*) was biotinylated using the Avidity BirA500 biotinylation kit (Avidity, EC 6.3.4.15) following the manufacturer’s instructions. rTTHC (Fina Biosolutions) and CRM197 (Fina Biosolutions) were biotinylated using Pierce EZ-Link biotinylation kit (Thermo Scientific, 31497). Excess biotin was removed by diafiltration with 100kDa cutoff. A 5-fold molar excess of biotin was used for the reaction.

### Flow cytometry

Briefly, LNs were collected into 1.5ml Eppendorf tubes and dissociated using a pestle. Single cell suspensions were incubated in Fc block (BD Biosciences) for 15min, followed by primary antibody staining at 4°C for 30min. For intracellular staining of Ki67, cells were permeabilized using BD Cytofix/Cytoperm kit, washed in the supplied permeabilization buffer and stained for 30min at 4°C.

Bait staining was performed with a combination of two different fluorescently-labelled streptavidins per bait. Biotinylated antigens were individually pre-incubated with each streptavidin-fluorophore before staining to allow tetramer formation, then cell suspensions were stained for 30min on ice, before staining for other surface markers as outlined above.

Antibodies used were as follows: Anti-mouse FC block (2.4G2, BD, 553142), Anti-mouse CD4 (RM4-5, Invitrogen, 47-0042-82), Anti-mouse CD95 (Jo2, BD, 557653), Anti-mouse CD38 (90, ThermoFisher, 56-0381-82), Anti-mouse B220 (RA3-6B2, BD, 748867), Anti-mouse CD138 (218-2, Biolegend, 142518), Anti-mouse TACI (8F10, BD, 742840), Anti-mouse CD86 (GL1, BD, 740688), Anti-mouse CXCR4 (2B11/CXCR4, BD, 558644), Anti-mouse CD45.1 (A20, Biolegend, 110708), Anti-mouse CD45.2 (104, Biolegend, 109839), Anti-mouse Ki67 (B56, BD, 561126), Anti-mouse TCRb (H57-597, Invitrogen, 47-5961-82).

### Cell sorting and VDJ Seq analysis

Single mouse GC B cells or PCs were purified and processed as described (*43*). Briefly, samples were single-cell sorted into 96-well plates containing 5ul lysis buffer (TCL buffer (Qiagen, 1031576); 1% 2-mercaptoethanol). Single cell Ig sequencing was performed using the nested PCR protocol (*43*). For IgL sequencing the following primers were used:

PCR1: 1mFL1 (ACTTATACTCTCTCTCCTGGCTCTC), 1mFL2 (CTCTTCTTCTTCTTTGTTCTTCATTGCT), 1mRL (GTACCATYTGCCTTCCAGKCCACT). Annealing temp=46°C. PCR2: 2mFL1 (CAGGCTGTTGTGACTCAG), 2mFL2 (CAACTTGTGCTCACTCAG), 2mRL (CTCYTCAGRGGAAGGTGGRAACA). Annealing temp=57°C. The 2mRL primer was used for sequencing. PCR products were Sanger sequenced and analyzed using Geneious Prime, and our Ig analysis bioinformatic pipeline (*44–46*). All scripts and data used to process antibody sequences are publicly available on GitHub at https://github.com/stratust/igpipeline/tree/igpipeline2_timepoint_v2. IgL, IgK and IgG/IgM IgH chains were paired (1xIgH, 1xIgL only) and analyzed.

### qPCR

200-500 cells were sorted into lysis buffer supplied with the SuperScriptIV Single Cell/Low Input cDNA preamp kit (Thermo Scientific, 11752048). Samples were treated as per kit instructions, and 13x cycles of preamplification were performed. Taqman probes for GAPDH(Mm99999915_g1), HPRT(Mm00446968_m1), Nr4a1(Mm01300401_m1), Irf4(Mm00516431_m1), Myc(Mm00487804_m1), Tfap4(Mm00473137_m1) were used for this assay, in conjunction with Taqman fast advanced master mix (Applied Biosystems, 4444556). For data presentation, relative expression was calculated normalized to GAPDH.

### ELISA

For detection of NP-reactive serum IgG, 96-half-well plates (Corning, 3690) were coated with 25ul NP_7_-BSA or NP_28_-BSA (Biosearch Technologies) at 10ug/ml in PBS overnight at 4°C. Plates were washed 2x with washing buffer (PBS, 0.05% Tween20, Sigma-Aldrich) and blocked in 150ul PBS, 2% BSA for 1h at RT. Plates were washed 4x more, and incubated for 2h at room temperature with serially diluted serum samples. B1-8^hi^ IgG was used as a standard on each plate to calculate absolute concentrations of NP_7_- and NP_28_-reactive IgG in serum. After 4 more washes, secondary HRP-conjugated anti-mouse IgG (Jackson, 115-035-071; 1:5000 dilution) was added for 1h at room temperature. Plates were washed 6x and developed using 3,3’,5,5’-tetramethylbenzidine (Thermo Scientific) for 3min, followed by addition of H_2_SO_4_. 450nm absorbance values were immediately measured using a microplate reader (Fluostar Omega, BMG Labtech).

### Confocal microscopy and analysis

Lymph nodes were prepared for confocal imaging as previously described (*47*). Briefly, LNs were collected from mice and fixed in 4% PFA for 2h. LNs were washed in PBS, immersed in 30% sucrose for cryoprotection for 12h, and embedded in OCT (Optimal cutting temperature compound; TissueTek) blocks. Tissues were sectioned at 7um on a cryostat (Microm) and blocked with mouse serum (Rockland) and Fc block for 30min. Slides were incubated at RT with the indicated antibodies for 6h, washed in PBS and mounted using fluoromount-G (ThermoScientific). Microscopy was performed on an inverted laser scanning microscope LSM980 (Zeiss).

For quantification, the Spots function in Imaris imaging software (Bitplane) was used. CD138^+^ PC were identified based on CD138-PE fluorescence intensity, cell quality (a metric which is automatically quantified in the Spots function), and these cells were sub-grouped based on staining intensity of NP and/or Ki67. All spots were manually verified.

### Single-cell libraries processing

For single-cell B cell receptor sequence reconstruction we used Cell Ranger (v.7.1.0) with mouse reference genome GRCm38. Hashtag-oligos (HTOs) UMI counts were processed using CITE-Seq-Count (v.1.4.5). We utilized MULTIseqDemux from Seurat (v4.1.1) to demultiplex, visualize and categorize cells as either GC or PC, based on the measured surface expression of CD138, CD95, CCR6 and CD86 by antibody-conjugated oligos. A small fraction of ZSG^+^ cells showing expression of CCR6 but lacking CD95 were included, but these cells were not classified as GC or PC, and were excluded from subsequent analysis.

### Computational analyses of antibody sequences

Single-cells heavy and light chains were paired using in-house R scripts and subsequently analyzed using igpipeline v2.0 (https://github.com/stratust/igpipeline/tree/igpipeline2_timepoint_v2), as previously described (*44*), using the mouse IMGT database (*48*) as reference. The paired IgH and IgL chains of antibodies from the same clonal progeny were merged and aligned to the mouse IMGT germline (GL) sequence using mafft v7.520 (*49*) with default parameters, except for –globalpair. Genotype-collapsed phylogenetic trees (GCtrees) of clonal lineages were inferred using GCTree v4.1.2 (https://github.com/matsengrp/gctree) (*50*). Each node represents a unique IgH and IgL combination with node size indicating the number of identical sequences. The small dotted nodes represent the unobserved ancestral. A CITE-Seq library including CD138, CD95, CCR6 and CD86 was used to assign GC or PC-identity to cell barcodes, and for UMAP generation of analyzed populations.

### Statistical analysis

Details of statistics including tests used, exact values and n numbers are indicated in figure legends and/or main text. Quantification and statistical analyses were performed in GraphPad Prism (Version 10.2.3), unless otherwise detailed in this methods section. Graphs generated using Prism were assembled into figures using Adobe Illustrator. Flow cytometry analysis was performed in FlowJo v.10.10.0 software (BD).

---

**Supplementary Fig. 1.**
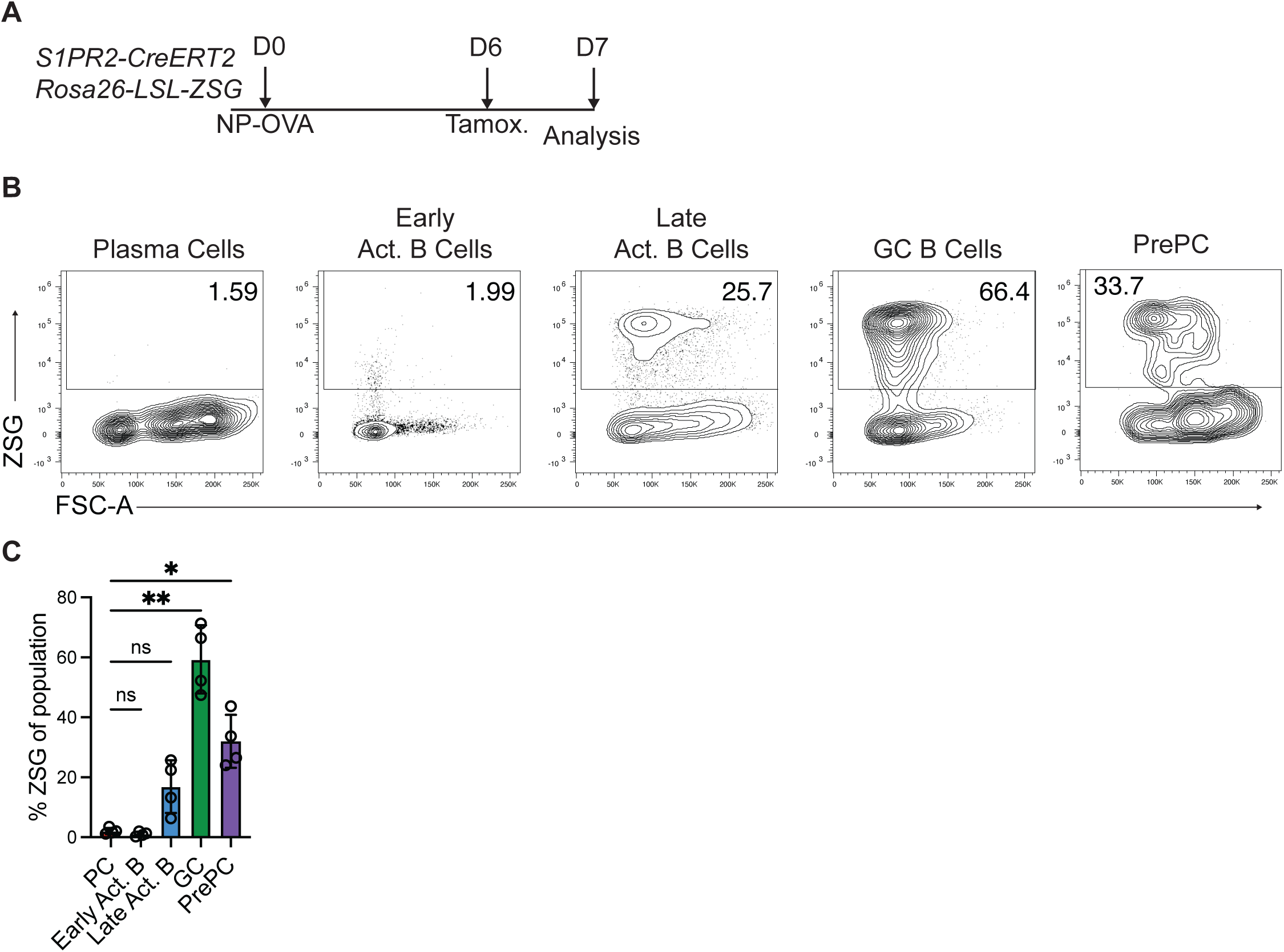
S1PR2-CreERT2 labels GC B cells and late activated B but not mature plasma cells, related to fig.1. (**A**) Experimental outline. (**B**) Representative flow cytometry plots showing ZSG expression in PCs (TACI^+^ CD138^+^), early activated B cells (TACI^-^ CD138^-^ B220^+^ CD38^+^ GL7^+^ Fas^-^), late activated B cells (TACI^-^ CD138^-^ B220^+^ CD38^+^ GL7^+^ Fas^+^), GC B cells (TACI^-^ CD138^-^ B220^+^ CD38^-^ GL7^+^ Fas^+^) and PrePC (TACI^-^ B220^+^ CD38^-^ GL7^+^ Fas^+^ CD138^+^). (**C**) Quantitation of data presented in B. Each point represents one mouse. Data in B-C represent one of three experiments performed. ns, not significant, ** P<0.005; ordinary one-way ANOVA.

**Supplementary Fig. 2.**
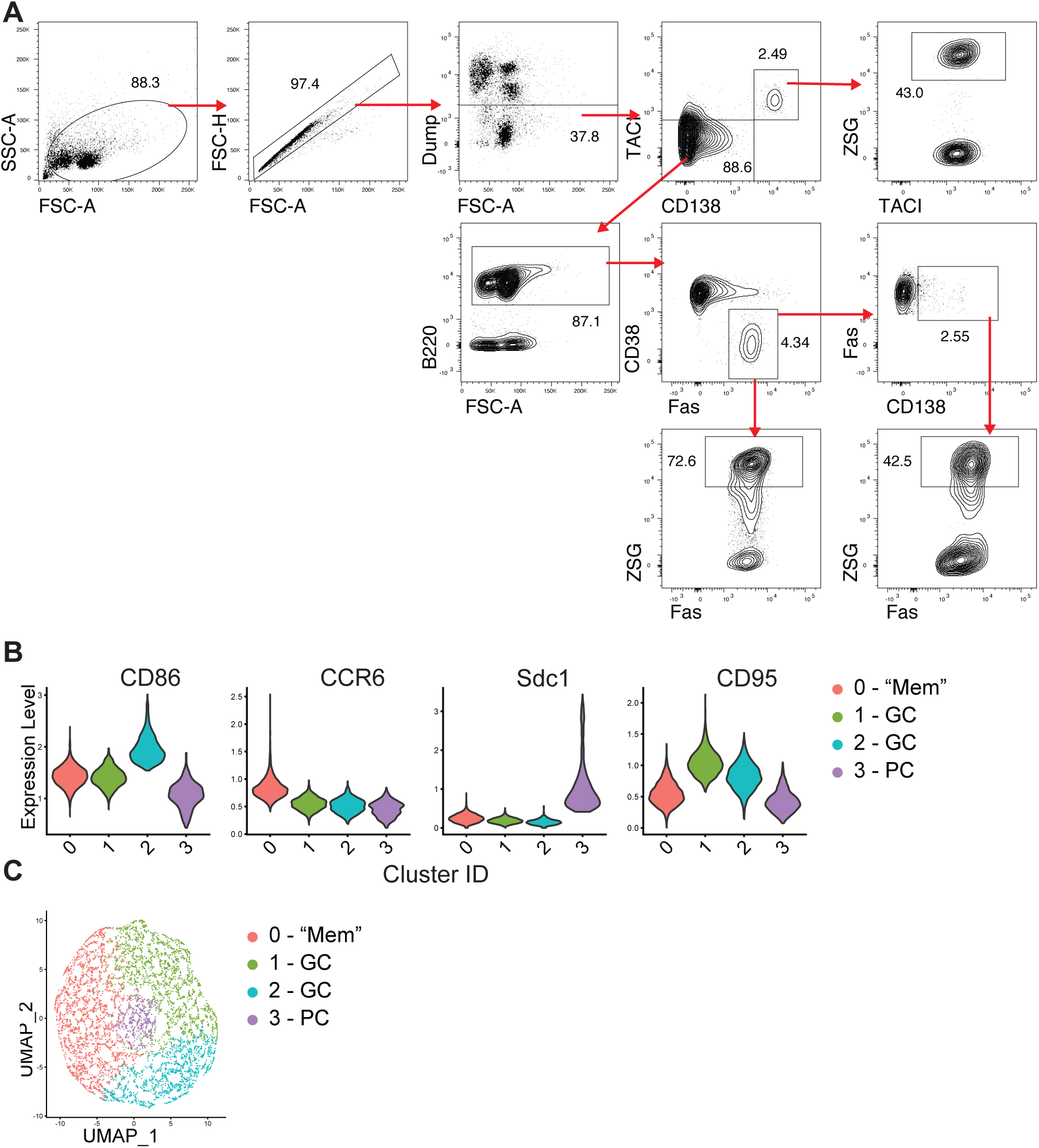
Identification of GC B and PC, related to fig.1. (**A**) Gating strategy for ZSG^+^ PCs (Dump^-^ CD138^+^ TACI^+^), GC B (Dump^-^ TACI^-^ B220^+^ CD38^lo^ Fas^+^) and prePCs (Dump^-^ TACI^-^ B220^+^ CD38^lo^ Fas^+^ CD138^+^), from animals treated as in Fig. 1A. Gating approach depicted was used for these populations throughout unless otherwise stated. (**B**) Expression levels of CITE-seq surface staining in clusters identified as PC and GC in two separate sequencing runs. (**C**) Uniform manifold approximation and projection (UMAP) of the above data. Data are pooled from 5 mice, and represent two experiments performed.

**Supplementary Fig. 3.**
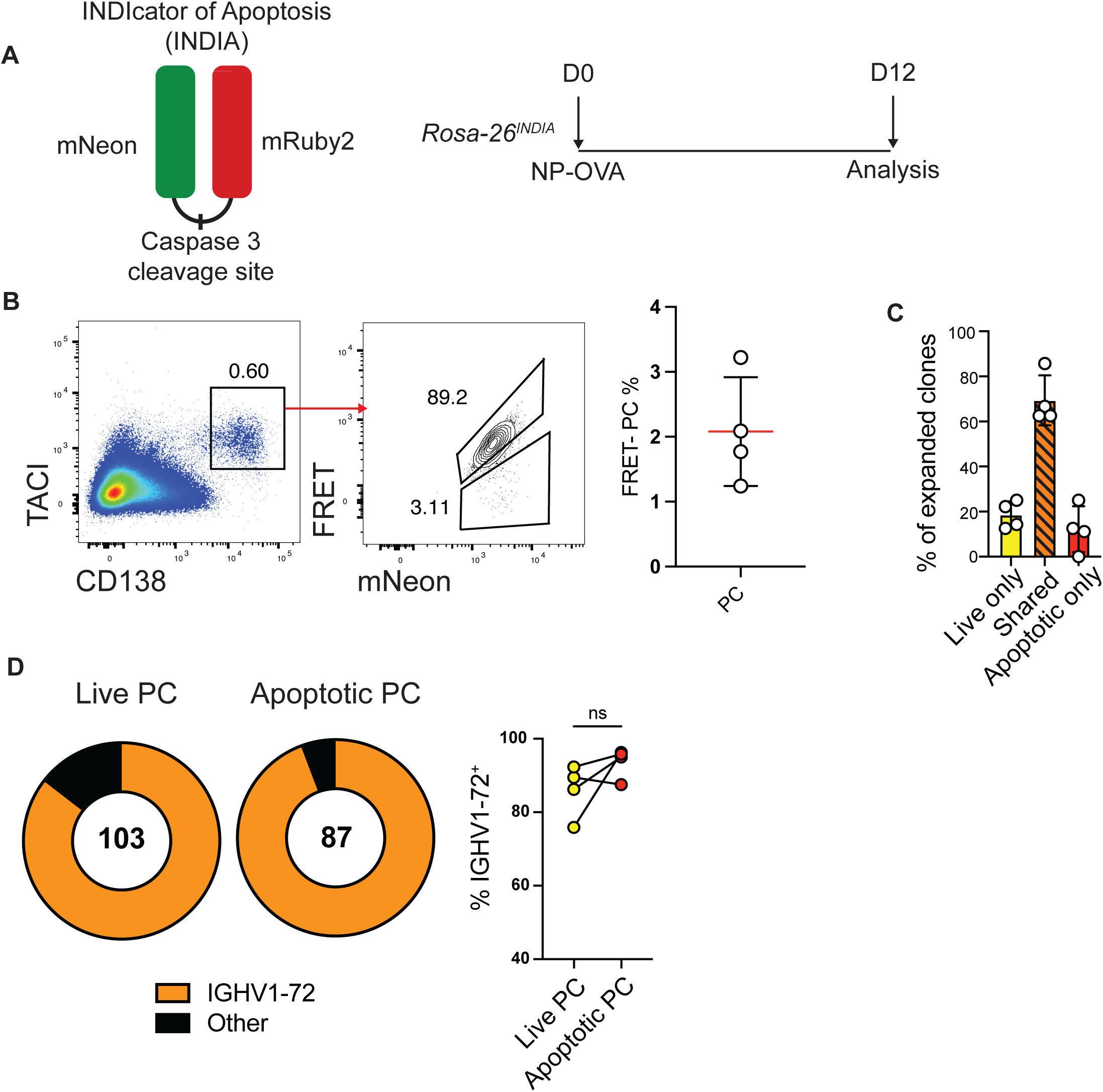
Plasma cell apoptosis is not associated with BCR affinity. (**A**) Left, Diagram of INDIA reporter. Right, Experimental outline for (**B-D**). (**B**) Left, representative flow cytometry plot showing gating for CD138^+^TACI^+^ PCs, FRET (BB630 channel) and mNeon. Right, Quantitation of FRET^+^ PCs. (**C**) Frequency of clones found only among live PCs, only in FRET^-^ apoptotic PCs or ‘shared’ clones found in both populations. Each point represents one mouse. (**D**) Left, frequency of FRET^+^ live PCs or FRET^-^ apoptotic PCs expressing IGHV1-72 antibodies. Right, summary of IGHV1-72 frequencies, each point represents one mouse. All experiments were performed at least twice.

**Supplementary Fig. 4.**
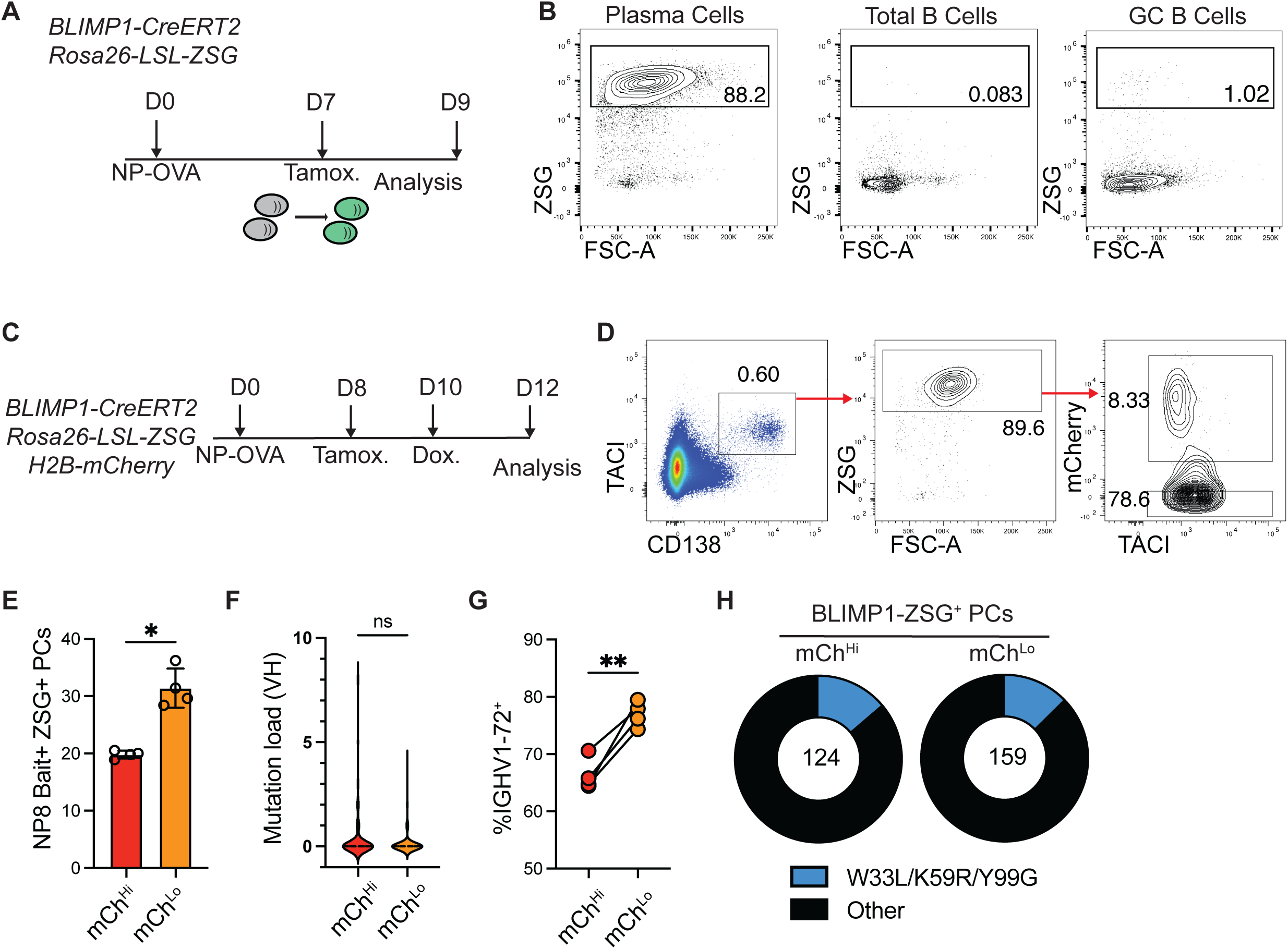
Proliferating PCs are enriched among high-affinity antigen binding cells. (**A**) Experimental layout for **B**. (**B**) Representative flow cytometry plots showing Blimp1-CreERT2-driven fate mapping of PCs (CD138^+^TACI^+^), total B cells (TACI^-^ CD138^-^ B220^+^) and GC B cells (TACI^-^ CD138^-^ B220^+^CD38^-^Fas^+^). (**C**) Experimental layout for **D-H**. (**D**) Flow cytometry profile showing TACI^+^CD138^+^ZSG^+^ PCs and gating for mCh^hi^ and mCh^lo^ cells from pLNs. (**E**) Quantitation of NP bait staining frequency among mCh^hi^ and mCh^lo^ PCs on D12 after immunization. (**F**) Number of VH mutations in ZSG^+^ PC populations.(**G**) Frequency of mCh^Hi^ or mCh^Lo^ ZSG^+^ PCs expressing IGHV1-72. (**H**) Frequency of high affinity mutation containing sequences among IGHV1-72^+^ expressing mCh^Hi^ or mCh^Lo^ ZSG^+^ PCs. ns, not significant * p<0.05, ** p<0.005. (**E, G**) paired two-tailed Student’s t-test; (**F**) unpaired Student’s t-test. Data are pooled from 2 independent experiments, n=4.

**Supplementary Fig. 5.**
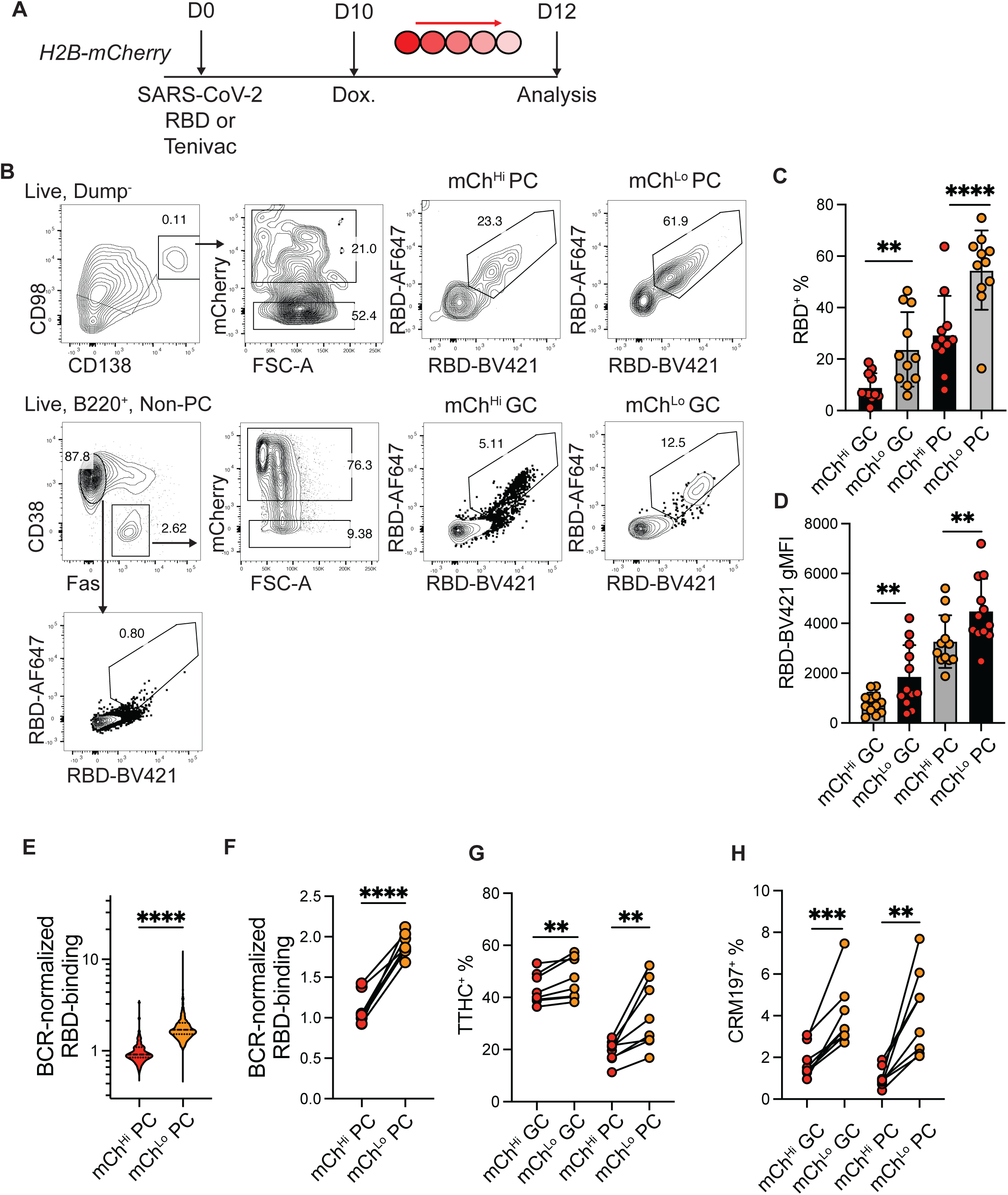
Proliferating PCs are enriched among high-affinity antigen binding cells. (**A**) Experimental layout used in (**B-D**). (**B**) Representative flow cytometry plots showing gating on mCh^hi^ and mCh^lo^ CD98^+^CD138^+^ PCs, and mCh^hi/lo^ B220^+^CD38^-^Fas^+^ GC B cells. Right panels display representative dual antigen bait staining for Sars-CoV-2 RBD. Naïve B220^+^CD38^+^Fas^-^ B cells were used as a negative control for bait staining. (**C**) Quantitation of RBD staining frequency among mCh^hi^ and mCh^lo^ GC B cells and PCs on D12 after SARS-CoV-2 RBD immunization. (**D**) Geometric mean fluorescence intensity (gMFI) of RBD-BV421 staining among mCh^hi^ and mCh^lo^ PC and GC B. (**E**) Violin plots displaying cellular distribution of BCR-normalized bait binding. (**F**) Average BCR-normalized bait binding. Each point represents one mouse. (**G-H**) Quantitation of tetanus toxoid heavy chain fragment c (TTHC; **E**) and detoxified diphtheria toxin (CRM197; **F**) bait binding on D12 after Tenivac immunization. * p<0.05, ** p<0.005, ***p<0.0005, **** p<0.0001. (**C-D, F-H**) Paired two-tailed Student’s t-tests. (**E**), Mann-Whitney test. Data in B-F and G-H are each representative of 3 independent experiments.

**Supplementary Fig. 6.**
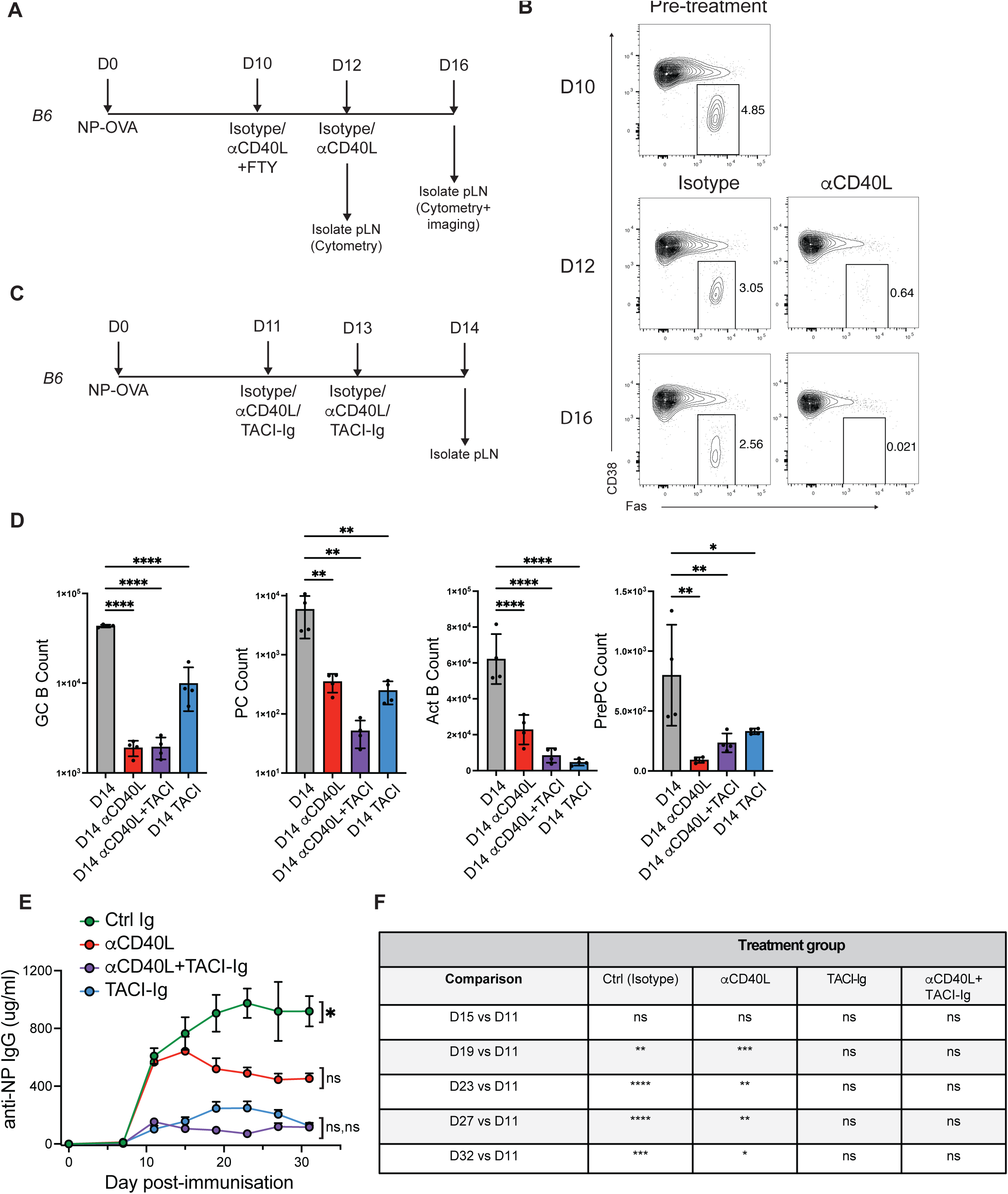
related to Fig. 3 and Fig.4. GC B and PC depletion kinetics. (**A**) Experimental layout used in (**B**). (**B**) Representative cytometry plots showing GC B cell depletion in pLNs on D12 and D16 after immunization. (**C)** Experimental layout used in (**D**). (**D**) Quantitation of GC B, PC, activated B and prePCs after treatment with depleting antibodies as described in (**C)**. (**E**) Total serum NP-binding IgG, from mice treated as in Fig. 4H, as measured by NP_28_-binding. Statistical comparisons shown represent results of a mixed-effects analysis, from endpoint D32 vs D11 onset of treatment. (**F**) Results of mixed effects analysis comparing affinity maturation (NP_7_/NP_28_ ratio) of the specified timepoints vs D11, in the same group. Data are presented in Fig.4D,E. * p<0.05, ** p<0.005, ***p<0.0005, **** p<0.0001, ns not significant. (**D**) Ordinary one-way ANOVA (all plots); (**E,F**) mixed-effects analysis.

**Supplementary Fig 7.**
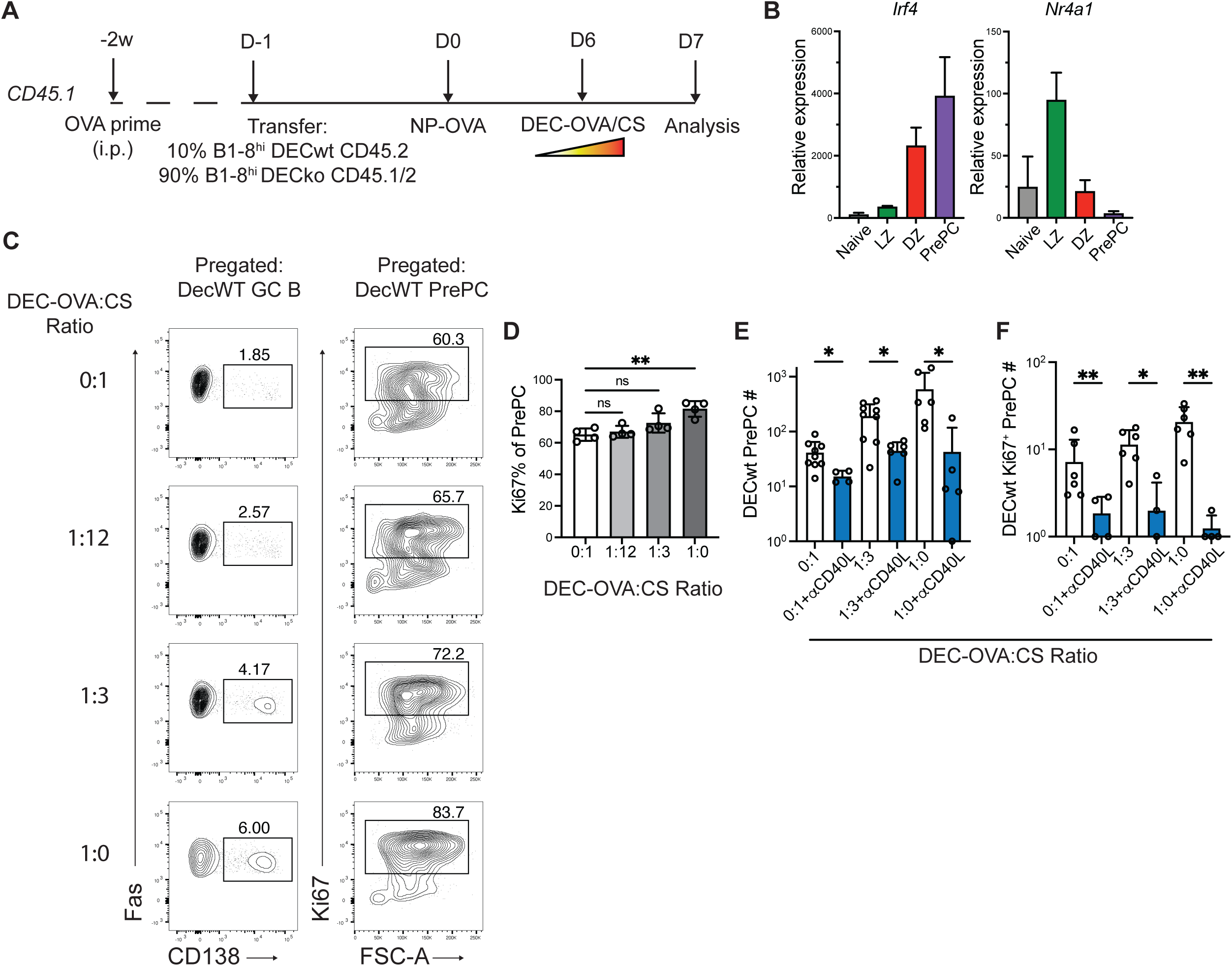
PrePC response to T cell help. (**A**) Experimental layout used in (**B-D**). (**B**) qPCR of purified naïve B (grey bars) LZ B (green bars), DZ B (red bars) and prePC (purple bars) showing GAPDH-normalized relative expression for *Irf4* (left) and *Nr4a1* (right). (**C**) Representative cytometry plots showing frequency of prePC and Ki67 staining among prePC, after DEC-OVA:DEC-CS administration. (**D**) Percentage of Ki67^+^ cells among total DEC^WT^ prePC. (**E-F**) Quantitation of DEC^WT^ prePCs (**E**) and Ki67^+^ DEC^WT^ prePCs (**F**) 72h after anti-DEC administration, with or without aCD40L treatment as indicated (also see Fig.5G). * p<0.05, ** p<0.005. (**D-F**) Kruskal-Wallis tests.

**Supplementary Fig 8.**
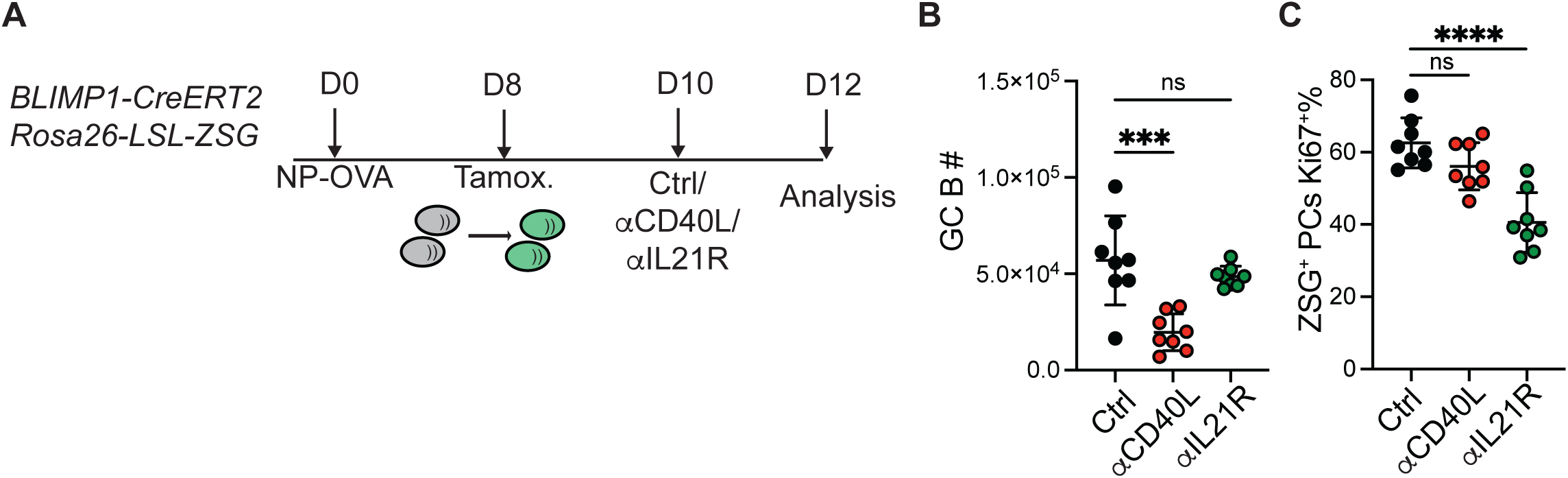
IL-21R supports post-GC expansion of PCs. (**A**) Experimental layout. (B) GC B cell numbers from mice treated with aCD40L, aIL-21R or isotype control antibodies between D10-D12. (**C**) Frequency of Ki67+ dividing cells amongst CD138^+^ TACI^+^ ZSG^+^ fate-mapped PCs. ns, not significant ** p<0.005; (**B**,**C**) Ordinary one-way ANOVA. Data are pooled from 3 independent experiments.

